# STX5’s flexibility in SNARE pairing supports Golgi functions

**DOI:** 10.1101/2022.05.24.493304

**Authors:** Zinia D’Souza, Irina Pokrovskaya, Vladimir V. Lupashin

## Abstract

The intracellular transport system is an evolutionally conserved, essential, and highly regulated network of organelles and transport vesicles that traffic protein and lipid cargoes within the cell. The events of vesicle formation, budding and fusion are orchestrated by the trafficking machinery – an elaborate set of proteins including small GTPases, vesicular coats, tethers, and SNAREs. The Golgi - the central organelle in this transport network, receives, modifies and sorts secretory and endocytic cargo. Glycosylation is one of the major modifications that occur within the Golgi, which houses enzymes and other components of glycosylation machinery. According to the current Golgi maturation model, Golgi resident proteins are constantly recycled from the late (trans) Golgi compartments to the early compartment (cis) by the evolutionary conserved vesicular trafficking machinery. The key modulator of vesicular trafficking and glycosylation at the Golgi is the Conserved Oligomeric Golgi (COG) complex – its interaction vesicular trafficking machinery particularly Golgi SNAREs (STX5, GS28 (GOSR1), GS15 (BET1L) and YKT6) that drive fusion of incoming vesicles. Since the COG complex functions upstream of SNARE-mediated vesicle fusion, we hypothesize that depletion of Golgi v-SNAREs would mirror defects observed in COG deficient cells. To test this, we created single and double knockouts (KO) of GS28 and GS15 in HEK293T cells and analyzed resulting mutants using a comprehensive set of biochemical, mass-spectrometry (MS) and microscopy approaches. Deletion of GS28 significantly affected GS15, but not the other two partners, STX5 and YKT6. Surprisingly, our analysis revealed that COG dysfunction is more deleterious for Golgi function than disrupting the canonical Golgi SNARE complex. Quantitative MS analysis of STX5-interacting SNAREs revealed unexpected flexibility of Golgi SNARE pairing in mammalian cells. We uncovered two novel non-canonical Golgi SNARE complexes – STX5/VTI1B/GS15/YKT6 and STX5/SNAP29/VAMP7 which were upregulated in GS28 KO cells. Analysis of cells co-depleted for GS28/SNAP29 or GS28/VTI1B SNAREs revealed escalated defects in Golgi glycosylation, indicating that upregulation of these complexes functionally substitutes deleted GS28. Our data points to the remarkable plasticity in the intra-Golgi membrane fusion machinery which is controlled by the COG complex.

## Introduction

Being highly dynamic and continuously delivered with cargo molecules bidirectionally, the integrity of the Golgi apparatus is key to intracellular membrane transport. Cisternal maturation, the currently accepted model for Golgi trafficking posits that anterograde cargo, delivered to cis-Golgi, is carried forward by gradual maturation of cis-Golgi to trans Golgi network (TGN), without leaving the Golgi (Glick and Luini, 2011; Kurokawa et al., 2019). As the cisternae mature, its resident proteins and enzymes are recycled back by vesicular retrograde trafficking. Golgi’s trafficking machinery includes small GTPases, coat proteins, cargo adaptors, tethers and SNAREs that work in concert to achieve its functions. The COG (Conserved Oligomeric Golgi) – the major multi-subunit tether (MTC (Multisubunit Tethering Complex) at the Golgi interacts with members from all classes of the Golgi vesicle trafficking machinery and plays a critical role in intra-Golgi trafficking (Blackburn et al., 2019; Shestakova et al., 2006; Willett et al., 2013b)

Vesicles recycling glycosylation enzymes are constantly fusing at the Golgi cisternae by SNARE-mediated fusion (Shestakova et al., 2006). SNAREs on the Golgi and vesicele membrane and form a low energy highly stable complex to provide the force for membrane fusion. In vitro studies using purified SNAREs on liposomes have shown that an appropriate set of SNAREs alone is sufficient to drive membrane fusion (McNew et al., 2000; Parlati et al., 2002). SNARE complex formation is highly specific and a fusogenic SNARE complex consists of 4 SNARE domains which are Qa, Qb, Qc and R (for review see (Jahn and Scheller, 2006)). These SNARE domains are provided by 3 or 4 SNARE proteins on the target and donor membranes. The major SNARE complex that controls intra-Golgi trafficking is STX5/GS28/GS15/Ykt6 (Xu et al., 2002), while the STX5/GS27/BET1/SEC22b operates in ER-Golgi anterograde cargo delivery (Xu et al., 2000) and STX16/VTI1A/STX6/VAMP4 regulates fusion of endosome-derived vesicles with TGN (Mallard et al., 2002). The participation of STX5/GS28/GS15/Ykt6 in a fusogenic SNARE complex, at the Golgi, was first reported by Yue Xu et al. (Xu et al., 2002). Immunoprecipitation with antibodies against STX5, GS28/GOSR1, GS15/BET1L and Ykt6 co-IP their other three SNARE partners (Xu et al., 2002; Zhang and Hong, 2001). Initial studies indicated that GS15 is localized in the cis-medial Golgi compartment (Xu et al., 2002), although a later immuno-EM study from Rothman lab observed that GS15 is present in a gradient of increasing concentration toward the trans side of the Golgi stack (Volchuk et al., 2004). Evidence that GS15 functions as a v-SNARE comes from the dominant-negative effect soluble GS15 has, when overexpressed (Xu et al., 2002). Soluble GS28 lacking its transmembrane domain was found to be membrane-bound leading to the speculation that GS28 is a t-SNARE by reasoning that GS28 lacking its transmembrane domain would only be membrane-bound by its association with its partners STX5 and Ykt6 (Rosenbaum et al., 2014). Both GS15 and GS28 are evolutionary conserved; their budding yeast homologs Sft1 and Gos1 localize and function at the Golgi (Bui et al., 1999; McNew et al., 1998; Xu et al., 2002; Xu et al., 1997). In vitro fusion studies with purified yeast SNAREs showed fusion was efficient with Sed5 (STX5 homologue), Gos1 and Ykt6 as t-SNAREs on acceptor liposomes and Sft1 as a v-SNARE on donor liposomes (Parlati et al., 2002).

COG-SNARE interactions, both in vivo and in vitro, have been well established by several studies (Kudlyk et al., 2013; Laufman et al., 2009; Sohda et al., 2010; Suvorova et al., 2002; Willett et al., 2013a; Zolov and Lupashin, 2005). The COG complex is predicted to tether COPI-formed intra-Golgi carriers (Cottam et al., 2014; Shestakova et al., 2006) and both GS28 and GS15 are present in COPI vesicles (Fusella et al., 2013; Gilchrist et al., 2006; Xu et al., 2002). GS28 and GS15 were described as “GEARs”-Golgi integral membrane proteins whose steady-state levels are decreased upon COG-subunit deletions/mutations. However, the abundance of STX5 and YKT6 is not regulated by COG (Oka et al., 2004). Furthermore, acute depletion of COG3 by siRNA-mediated knockdown (KD) results in the accumulation of small trafficking intermediates termed COG-complex dependent (CCD) vesicles which contain GS28 and GS15 (Zolov and Lupashin, 2005). While GS15’s role as a v-SNARE is clear, it is possible that GS28 also functions as a v-SNARE and travels on COPI intra-Golgi vesicles together with GS15 to form a fusogenic SNARE complex with STX5 and Ykt6 during vesicle fusion at the Golgi. The COG complex not only maintains the steady-state abundance and Golgi localization of v-SNAREs but is also required for SNARE complex formation. Upon COG3 or COG7 KD in HeLa cells, the amount and/or stability of STX5 complex containing GS28 and GS15 is dramatically reduced (Shestakova et al., 2007). COG4 physically interacts with STX5 and STX5 partner hSly1/SCFD1 via its N-terminal domain (Laufman et al., 2009). Over-expression of a COG4 N-terminal 153aa-long fragment as well as expression of E53/71A COG4 mutant disrupted the Golgi morphology, mislocalized GS15 and dramatically decreased GS28/STX5 interaction (Laufman et al., 2013b; Laufman et al., 2009). Similarly, GS15 or STX5 KD in several cell lines resulted in Golgi fragmentation (Suga et al., 2005; Xu et al., 2002).

While the exact functional role of the COG complex in the Golgi physiology is yet to be fully understood, in the current model the COG complex is thought to orchestrate intra-Golgi trafficking by capturing vesicles and modulating/proof-reading the SNARE alignment that precedes fusion (Laufman et al., 2013b; Willett et al., 2013b). Therefore, we hypothesized that the impairment in Golgi SNARE interactions, the critical step for vesicle fusion at the Golgi, is the major contributor to glycosylation defects caused by COG malfunctions. Indeed, GS28 along with COG subunits are frequently identified as hits in CRISPR screens investigating Golgi key trafficking and glycosylation players (Diep et al., 2019; Morgens et al., 2017; Tian et al., 2018; Yamaji et al., 2019). We reasoned that knocking out either GS28 or GS15 would phenocopy COG KD/KO glycosylation defects because Golgi enzymes will be mislocalized to transport vesicles unable to fuse with the Golgi and subsequently be degraded in GS28/GS15-depleted cells. In this study, we successfully knocked out GS28 and GS15 in HEK293T cells and characterized trafficking and glycosylation defects associated with the deletion of Golgiv-SNAREs. Surprisingly, we discovered that both GS28 and GS15 are mostly dispensable for Golgi glycosylation. To accommodate STX5-mediated retrograde trafficking of Golgi enzymes, mutant cells compensate for the absence of Golgi v-SNAREs by repurposing several post-Golgi SNARE proteins, primarily SNAP29, VAMP7 and VTI1B.

## Results

### HEK293T cells lacking SNAREs GS28 and GS15 are viable and maintain normal Golgi morphology

Using a CRISPR-Cas9 approach, we successfully knocked out (KO) GS28, the Golgi Qb SNARE and GS15, the Qc SNARE from the STX5/GS28/GS15/Ykt6 SNARE complex. Individual HEK293T clones lacking the targeted SNAREs were selected by western blotting (WB) (Figure 1A). GS28 and GS15 are Golgi localized SNARES and immunofluorescence (IF) was used for further confirmation of KO (Figure 1B). The resultant GS28 and GS15 KO clones were viable and did not show either altered cell morphology or growth delays (Z.D. unpublished observation). Giantin/GOLGB1 and GM130/GOLGA2 staining showed that the cis-Golgi morphology was unaffected by the deletion of either GS28 or GS15. The Golgi continuity and size in the SNARE KO cells appeared similar to WT Golgi. We thus obtained HEK293T clones deficient for GS28 or GS15 and evaluated the effect of their depletion on Golgi physiology

**Figure 1:**
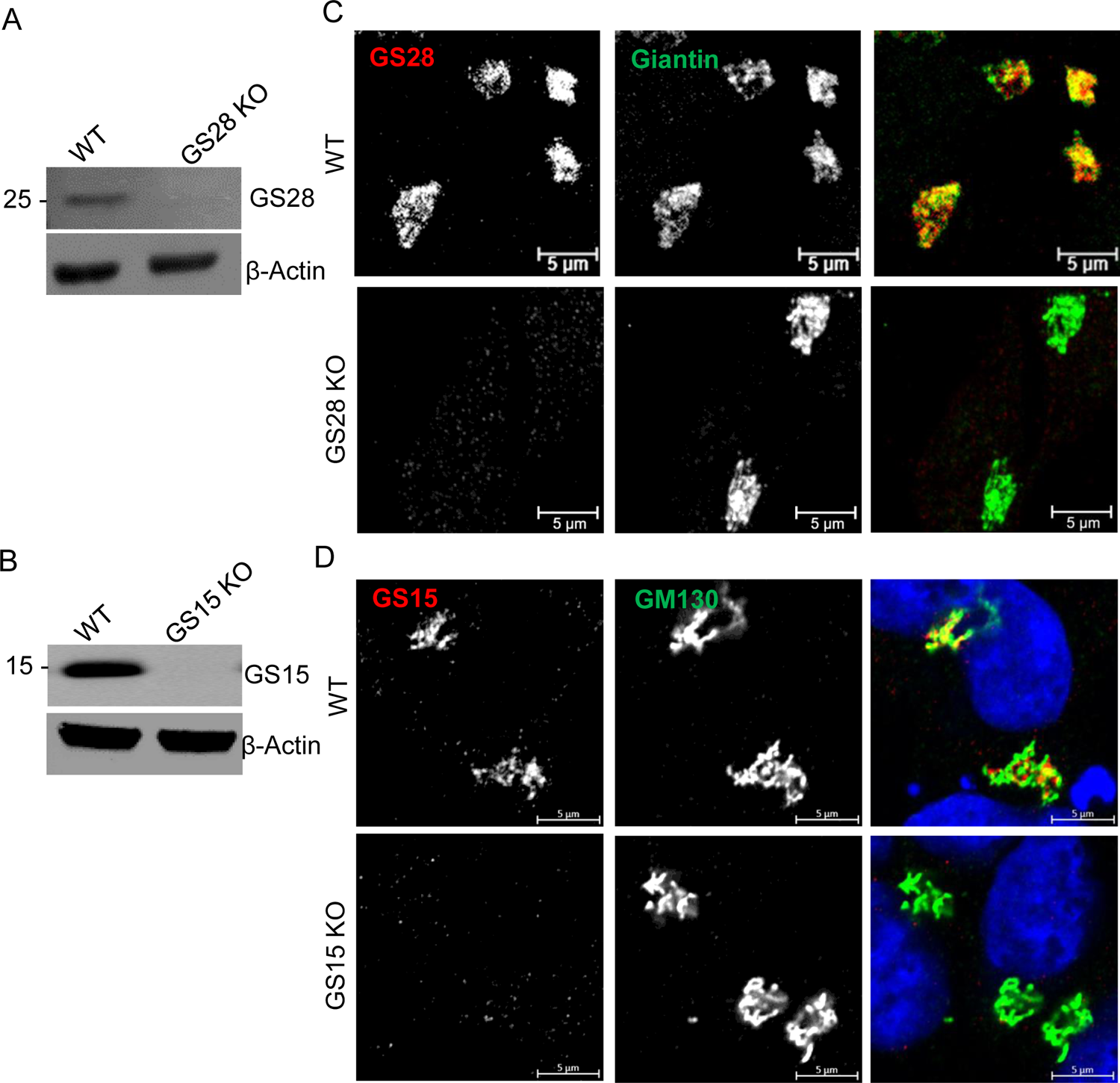
HEK293T cells lacking SNAREs GS28 and GS15 are viable and maintain normal Golgi morphology. (A, B) WB analysis of GS28 KO and GS15 KO in HEK293T-Cas9 cell line; (C, D) IF reveals complete loss of Golgi localized GS28 or GS15 in the KO cell lines. The Golgi is stained by Giantin and GM130 respectively. Scale bars are 5µm

### GS28 KO alters the stability and localization of GS15 but does not affect STX5 and YKT6

Thereon, we measured the protein levels of the STX5 SNARE partners using WB (Figure 2A, B). In GS28 and GS15 KO cells, the total protein levels of STX5 and YKT6 were unaffected. GS15 deletion did not affect the abundance of GS28. However, upon GS28 deletion, the level of GS15 was significantly decreased. IF also showed that the signal intensity of Golgi localized GS15 was significantly reduced in GS28 KO but GS15 KO did not affect of the Golgi localization of GS28 (Figure 2C, D). We have also created a GS28/GS15 double KO (DKO) HEK293T cells line and for all tested parameters, DKO behaved identically to GS28 KO (ZD, unpublished data). Comparing COG KO effects on the STX5 SNARE partners, GS28 KO partially phenocopied COG KO. In other words, GS28 and GS15 were significantly depleted in COG KOs similar to the GS28 KO phenotype. We then sought to address whether deleting the core fusion machinery at the Golgi will phenocopy any of the effects of COG malfunction.

**Figure 2:**
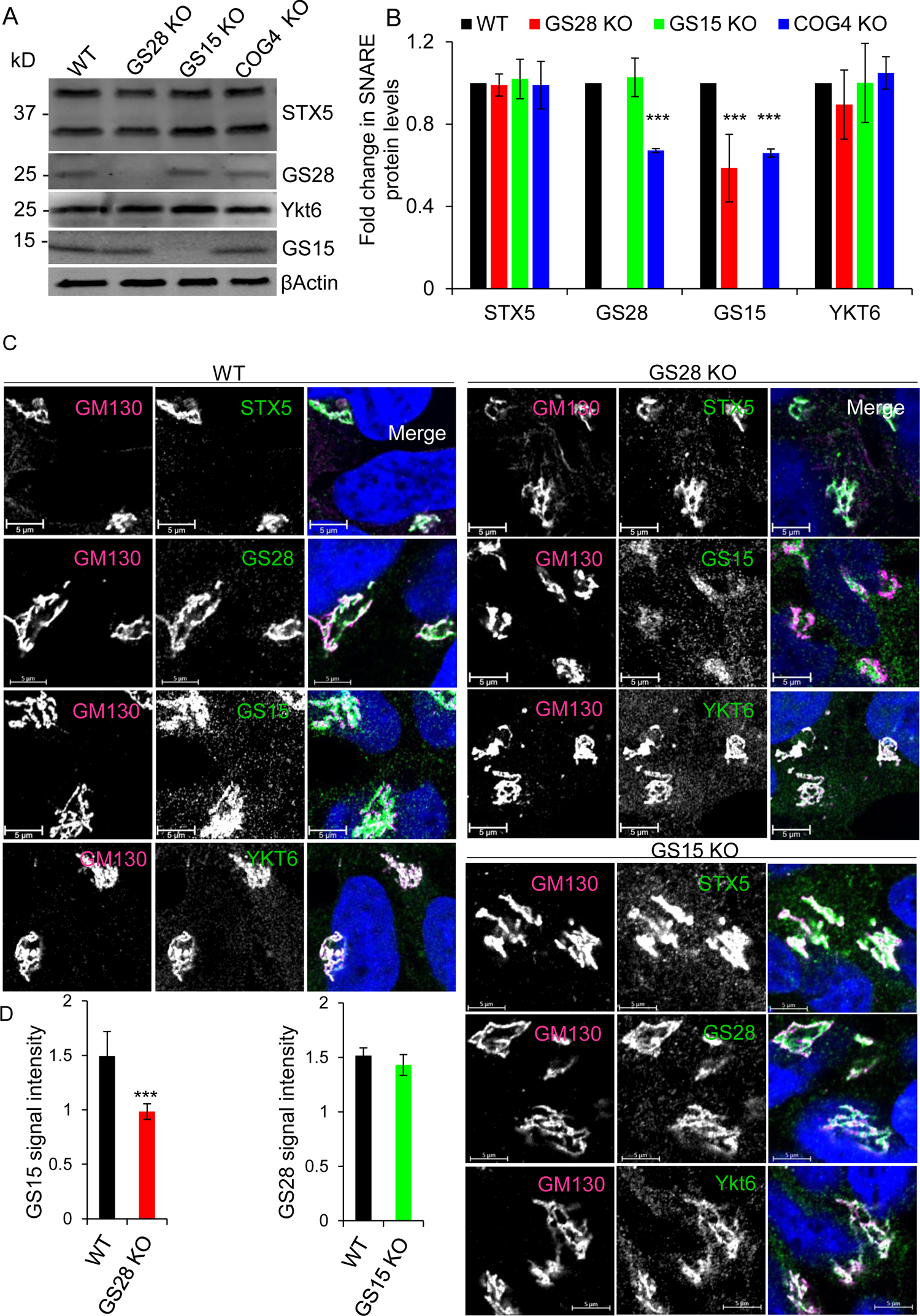
GS28 KO alters stability and localization of GS15 but does not affect STX5 and YKT6. (A, B) WB and quantification for the Golgi SNAREs in WT, GS28 KO, GS15 KO and COG4 KO cells showing significantly reduced levels of GS15 in GS28 KO cell lysates. In COG4 KO cell lysates, both GS28 and GS15 are significantly depleted. The amount of STX5 and YKT6 is unaffected by either SNARE or COG KO; (C) IF for STX5, GS28, GS15 and Ykt6 in WT, GS28 KO and GS15 KO cell lines shows all remaining STX5 partners are mostly Golgi localized in KO cells. The Golgi is stained with GM130. Scale bars are 5 µm; (D) The total intensity of Golgi localized GS15 is significantly reduced in GS28 KO but GS15 KO does not have any effect GS28’s intensity at the Golgi. Quantification of GS15’s and GS28’s signal intensities in the Golgi region in GS28 KOs and GS15 KOs respectively. n≥30, ***p>0.001

### Knocking out GS28 impairs retrograde trafficking

Subtilase cytotoxin (SubAB), an AB_5_ toxin exploits the cell’s retrograde trafficking pathway. It gets transported from the plasma membrane via Golgi to the ER where it cleaves an ER-resident protein, GRP78/BiP. Cells with defects in the retrograde trafficking machinery are protected from BiP cleavage by SubAB (Blackburn and Lupashin, 2016; Paton and Paton, 2010; Smith et al., 2009). Therefore, we used the SubAB assay to assess the integrity and proper function of Golgi retrograde trafficking in Golgi SNARE depleted cells. In WT cells, there was complete cleavage of GRP78 180min after the exposure of the cells to the toxin (Figure 3A, B). A similar dynamic of GRP78 cleavage was observed in GS15 KO cells, indicating that this SNARE is disposable for SubAB trafficking. However, in the case of the GS28 KO, only 50% of GRP78 was cleaved, similar to incomplete GRP78 cleavage observed in COG4 KO cells (Figure 3A, B, (Bailey Blackburn et al., 2016)). Altered SubAB trafficking may also indicate changes in Golgi integrity. As we have shown previously (Figure 1), GS28 and GS15 KOs did not change the integrity of the cis-Golgi compartment. To investigate whether there is any alteration in the overall Golgi morphology, we stained cells for cis (GM130) and trans (TGN46/TGOLN2) Golgi markers and measured their colocalization using super-resolution Airyscan microscopy. The analysis found a significant reduction in the colocalization between GM130 and TGN46 in the GS28 KO compared to WT. Furthermore, there was a reduced intensity of TGN46 staining in the GS28 KO compared to WT. In GS15 KO cells GM130-TGN46 colocalization was not significantly reduced, while in COG4 KO cells relative cis-trans Golgi colocalization was slightly increased, probably due to a severe fragmentation and intermixing of both compartments (Figure 3C, D, (Bailey Blackburn et al., 2016)). Taken together, our data indicateThe retention and recycling of enzymes that GS28 KO has impaired Golgi retrograde trafficking and/or overall Golgi integrity, although these changes differ from the extensive Golgi fragmentation observed in COG KO cells. Since the GS28 KO phenocopied the COG KO retrograde trafficking defect, we wondered whether Golgi glycosylation would also be affected in SNARE deficient cells.

**Figure 3:**
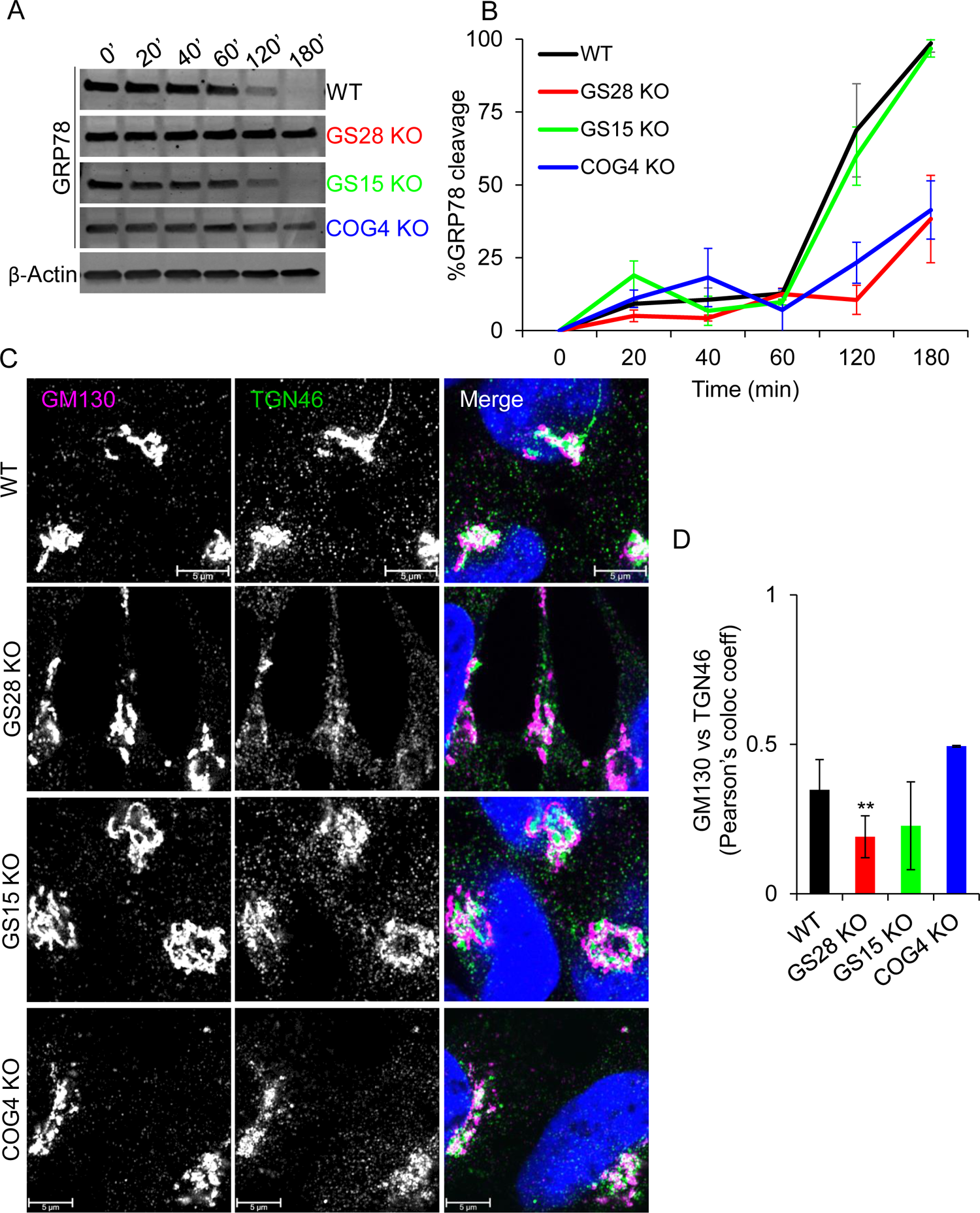
GS28 KO phenocopies COG4 KO retrograde trafficking defects. GS28KO alters endosome-to-Golgi delivery of SubAB toxin and TGN46. (A) WB showing GRP78 levels in lysates from WT, GS28 KO, GS15 KO and COG4 KO cells treated with SubAB toxin for indicated time periods. (B) Quantification from GRP78 levels (n=3). After 180 mins treatment with SubAB, 100% GRP78 is cleaved in WT and GS15 KO cells. However, in this time period only 50% is cleaved in GS28 KO and COG4 KO cells. (C) IF analysis of the recycling TGN46 in SNARE-depleted cells shows that TGN46 staining intensity in GS28 KO cells is reduced. (D) Pearson’s colocalization coefficient between GM130 and TGN46 **p<0.005, n≥30

### GS28 is essential for proper protein and lipid glycosylation

Glycosylation is a template-independent process which is entirely dependent on the sequence of enzyme-substrate interactions (D’Souza et al., 2021). Anterograde cargo molecules delivered from the ER, traverse the Golgi by cisternal maturation, and encounter glycosylation enzymes localized in each of the Golgi cisternae. Golgi is tasked with transporting cargo forward and ensuring its glycosylation machinery including glycosylation enzymes are properly compartmentalized. The retention and recycling of enzymes are intimately tied to the balance in anterograde and retrograde trafficking at the Golgi. Mutations or knockout of COG subunits severely impair Golgi retrograde trafficking. Acute knock-down (KD) or KO of any of the COG subunits results in mislocalization of every tested Golgi enzyme (Bailey Blackburn et al., 2016; D’Souza et al., 2019; Pokrovskaya et al., 2011; Shestakova et al., 2006; Zolov and Lupashin, 2005). We used WB to measure the total cellular level of four endogenous glycosylation enzymes involved in both N-(MGAT1 and B4GALT) and O-(GALNT2 and GALNT3) glycosylation. When we measured the protein levels of glycosyltransferases in control and SNARE KO cells, we found that the abundance of the tested enzymes was similar to control in GS15 KO cells and significantly reduced in GS28 KO, but this reduction was not as dramatic as in COG4 KO’s (Figure 4A, B). Since expression of GS28 was found to be required in maintaining the normal levels of B4GALT1, MGAT1 and GALNT3 we wondered whether partial depletion of Golgi enzymes would affect Golgi glycosylation in GS28 KO cells. First, we examined potential changes in electrophoretic mobility of Golgi-glycosylated LAMP2, SDF4 and TMEM165 (Fukuda, 1991; Scherer et al., 1996; Stribny et al., 2020). The predicted molecular weight of LAMP2 is about 45 kD. However, fully processed Lamp2 runs close to 100kD. In the COG4 KO cell lysate, there was a dramatic increase in the electrophoretic mobility of LAMP2 indicating severe glycosylation defects. In GS28 KO, but not in GS15 KO cells, there was a subtle increase in LAMP2’s electrophoretic mobility. GS28 KO did not show any effect on SDF4 but affected the electrophoretic mobility and stability of TMEM165 (Figure 4C, D). These results indicate that GS28 is needed for efficient Golgi glycosylation, but its deletion does not affect glycosylation as drastically as COG’s deletion.

**Figure 4:**
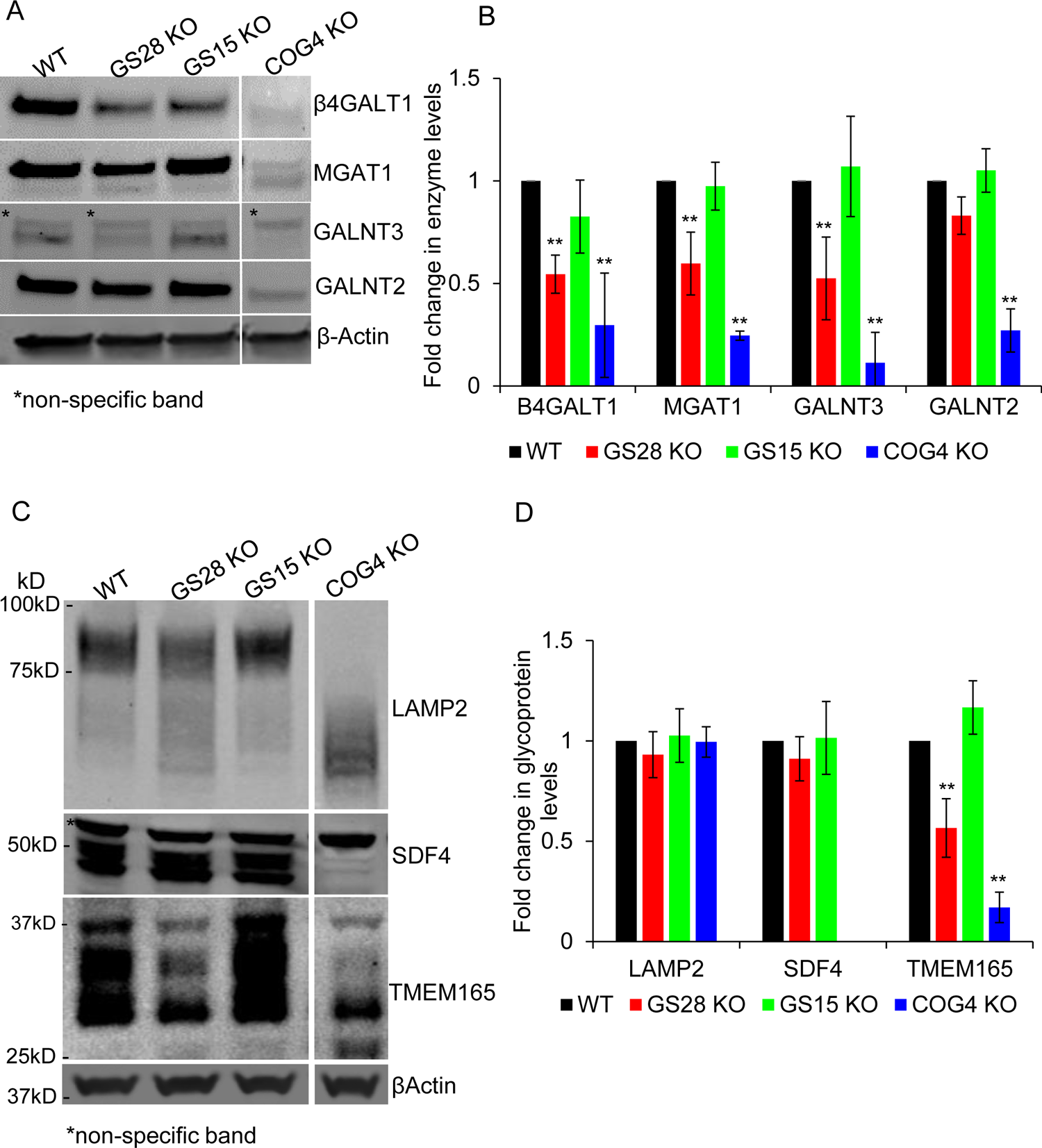
The abundance of several Golgi enzymes is reduced on GS28 KO cells but to a lesser degree as in COG depleted cells. Enzyme reduction in SNARE-depleted cells did not significantly alter glycosylation of lysosomal and Golgi **glycoproteins.** (A) WB analysis of Golgi glycosylation enzymes in WT, GS28 KO, GS15 KO and COG4 KO cells. While the total amount of B4GALT1, MGAT1, GALNT3 and GALNT2 is not as severely depleted in GS28 KOs compared to COG4 KOs, their levels are significantly reduced compared to the WT. (B) Quantification of fold changes in enzyme levels with respect to WT from biological triplicates. (C) WB showing electrophoretic mobility and abundance of glycoproteins; (E) Fold change in glycoprotein levels from biological triplicates with respect to WT. **p<0.001, ***p<0.0001

We further assessed glycosylation defects in SNARE KOs by utilizing fluorescently labelled lectins. GNL and HPA are lectins that recognize early N- and O-linked glycans. GNL binds high-mannose residues on N-glycans and HPA binds alpha N-acetylgalactosamine residues also known as core-2 structures on O-glycans (Brooks, 2000; Shibuya et al., 1988). Both these glycans are intermediates of the Golgi N- and O-glycosylation pathways. There was a significant increase in GNL-647 and HPA-647 binding to the total cellular glycoproteins isolated from GS28 KO cells (Figure 5A, B). This suggests an increase in the accumulation of immature glycans. However, compared to GS28 KO, COG KO resulted in more than a 5-fold greater increase of immature N- and O-linked glycans. Immunofluorescence and flow-cytometric measurement of fluorescent HPA or GNL binding to plasma membrane glycoconjugates showed the peaks shift to the right of the WT in the case of GS28 and COG4 KOs (Figure 5D, E). This result indicates that GS28 KO cells accumulated immature and incomplete N- and O-linked glycans exposed on their surface.

**Figure 5:**
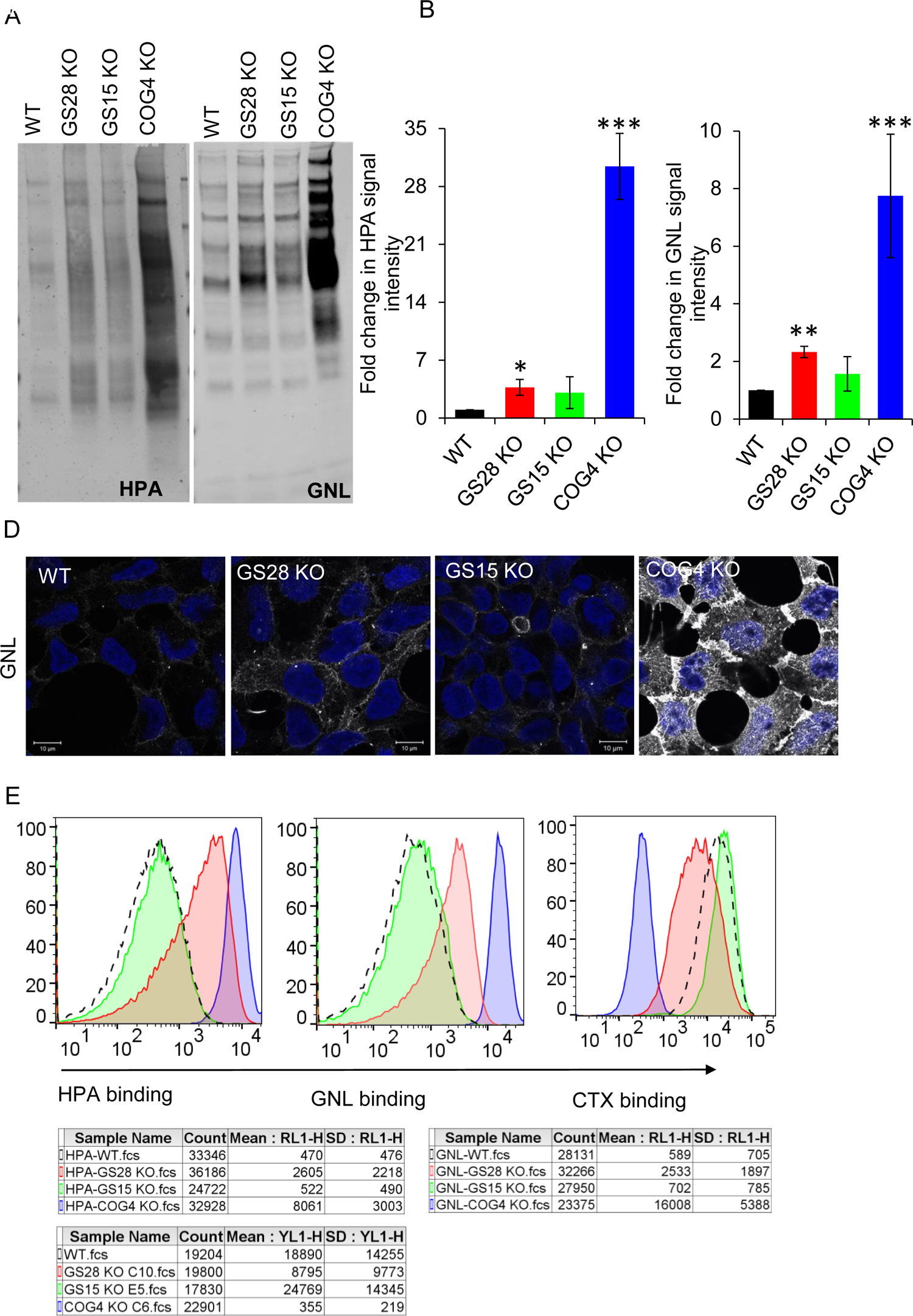
Lectin staining reveals mild glycosylation defects in GS28 cells. (A) Lectin blot detects the total amount of immature O-glycan - core-2 labelled by HPA-647, and N-glycan – high mannose labelled by GNL-647, in cell lysates (B) Quantification in the fold change in HPA and GNL staining intensity with respect to WT. There is a significant increase in the accumulation of immature glycoproteins in GS28 KOs but this is not as dramatic as in COG4 KOs. (C) Cell-surface labelling of immature N-glycan, high-mannose, stained with GNL-647 in WT and KO cells. (E) Flow-cytometry quantification of fluorescent HPA, GNL and CTX binding to the cell surface of wt and mutant cells. Incomplete glycan processing in the Golgi results in the expression of immature N and O-linked glycoproteins on the cell surface to which GNL and HPA bind, respectively. Reduced binding of CTX to GS28 KOs, which binds to GM1, a glycolipid indicates GS28 KO all affects glycolipid processing.

Cholera toxin (CTX), an AB_5_ toxin binds with high affinity to GM1 on the plasma membrane (Kenworthy et al., 2021). GM1 is a glycolipid, specifically, a monosialoganglioside. COG KO cells are resistant to CTX binding due to impaired lipid glycosylation (Bailey Blackburn et al., 2016). Flow-cytometric measurement of CTX-555 binding to the cell surface showed the peaks shift to the left of the WT in the case of GS28 and COG4 KO (Figure 5E, (Bailey Blackburn et al., 2016)). Like for the other two lectins, the decrease in CTX binding in GS28 KO cells was significantly smaller as compared to COG4 KO cells. Taken together, our glycosylation studies in SNARE KOs indicate that GS28, but not GS15 is needed for proper glycosylation but the effect of GS28 KO on Golgi glycosylation is significantly weaker compared to the effects observed in COG KO cells. We were surprised that deleting a key SNARE, GS28, from the Golgi fusion machinery, did not severely affect the maintenance of Golgi glycosylation machinery and Golgi glycosylation. Since in vitro binding studies have shown SNAREs are promiscuous in their binding preferences (Bethani et al., 2007; Tsui and Banfield, 2000), we entertained the possibility that in GS28 KO cells GS28 (and possibly GS15) is substituted by other SNAREs to accommodate sufficient vesicle recycling of Golgi resident proteins. To test this possibility, we looked for potential changes in STX5 partners in GS28 KO cells.

### GS28 KO impairs the partnering of the Golgi STX5 complex, and this leads to non-canonical SNARE substitution for missing GS28

Immunoprecipitation (IP) with affinity-purified antibodies against STX5 followed by quantitative label-free DIA analysis was performed to determine the effect of GS28 KO on STX5’s partnering with other SNARE proteins (Supplemental table 1). In this approach, cells were pretreated with NEM to inhibit the activity of NSF (Block et al., 1988; Rothman, 1994; Wilson et al., 1992) and thereby prevent dissociation of SNARE complexes during isolation. GS28 KO resulted in about 2-fold reduction of STX5 interactions with GS27/GOSR2, BET1 and GS15 (Figure 6A, B, C, supplemental table 1), indicating that GS28 KO not only impairs the Golgi STX5/GS28/GS15/Ykt6 SNARE complex but also affects the ER-Golgi STX5/GS27/BET1/Sec22b SNARE complex (Figure 6A, B). On the other hand, GS28 KO significantly increased STX5’s partnering with SNAP29, Vti1b and VAMP7 (Figure 6A, B, C, supplemental table 1). The unbiased mass-spectrometry (MS) results suggest that GS28 deletion is compensated for by the formation of one or more “non-canonical” STX5-containing complexes, which potentially operate at the Golgi under “SNARE-stress” conditions. It is likely that GS15 was also partially substituted since, in absence of GS28, the total levels of GS15 and the amount of GS15 coIPed with SXT5 are reduced.

**Figure 6:**
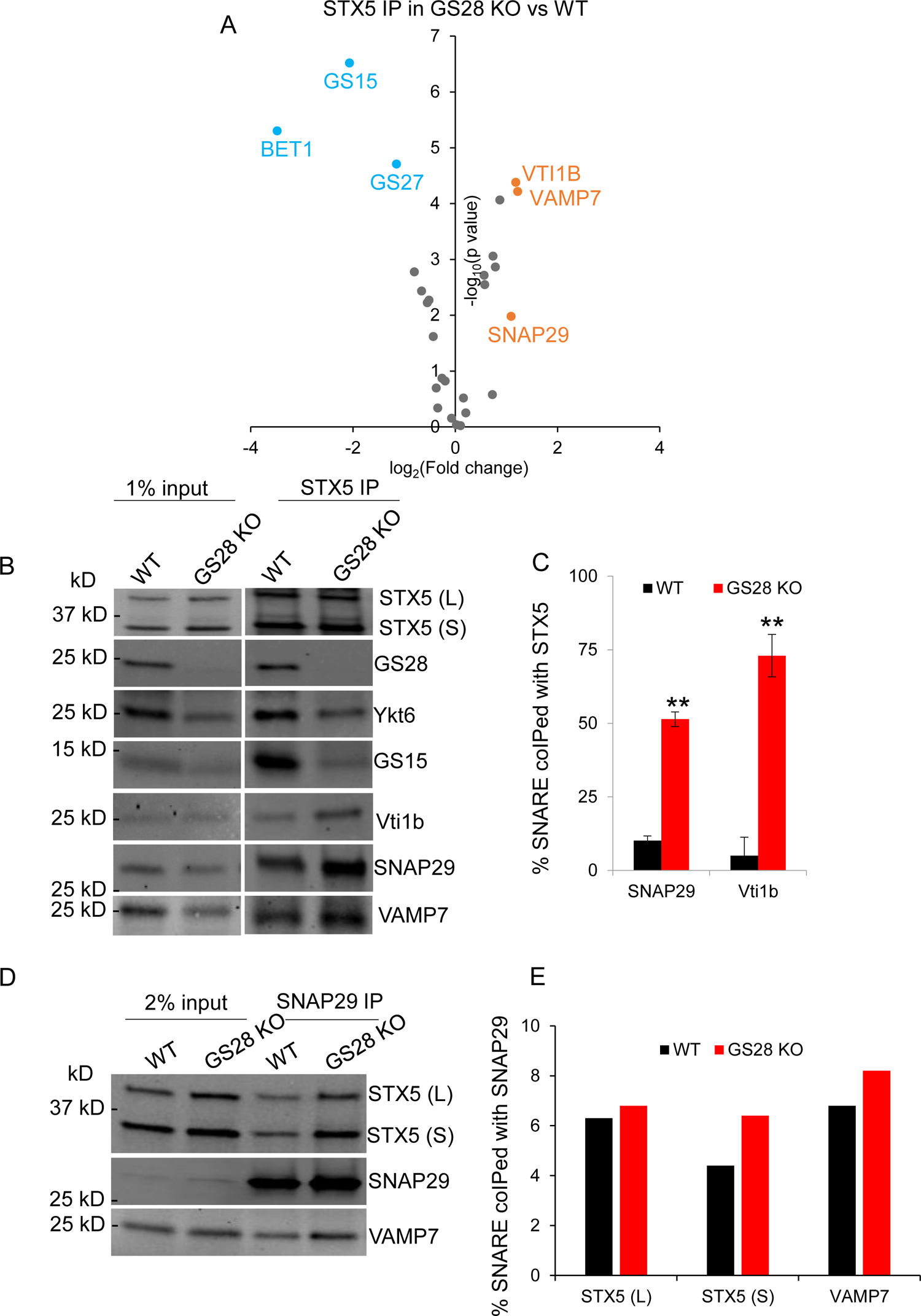
Label-free mass spectrometry (DIA MS) analysis of STX5 binding proteins reveals an increase in non-canonical STX5 SNARE partners. (A) Volcano plot showing fold change in SNAREs coimmunoprecipitated with STX5 in GS28 KO vs WT from DIA MS of four biological replicates. Red dots indicate significant increase and blue indicate significant decrease in GS28 KO vs WT. (B, C) WB analysis confirms MS results showing SNAP29 and Vti1b are significantly increased in STX5 IP from GS28 KOs. **p<0.001. (D, E) SNAP29 IP in WT and GS28 KO cells confirms the existence of STX5-SNAP29-VAMP7 SNARE complex.

We proposed that GS28 depletion stimulated two different “non-canonical” SNARE assemblies. Qb SNARE Vti1b may substitute for GS28 to form STX5/Vti1b/GS15/Ykt6 complex and Qbc SNARE SNAP29 may substitute both GS28 and GS15 to form STX5/SNAP29/R-SNARE complex. In the latter scenario, the canonical Golgi R-SNARE YKT6 is likely to be substituted with VAMP7, since the SNAP29 IP showed a significant preference for that R-SNARE as a partner (Figure 7, Supplemental figure 3). WB analysis of the STX5 IP suggests that both “non-canonical” complexes exist in cells under normal conditions and their abundance significantly increased upon GS28 deletion.

**Figure 7:**
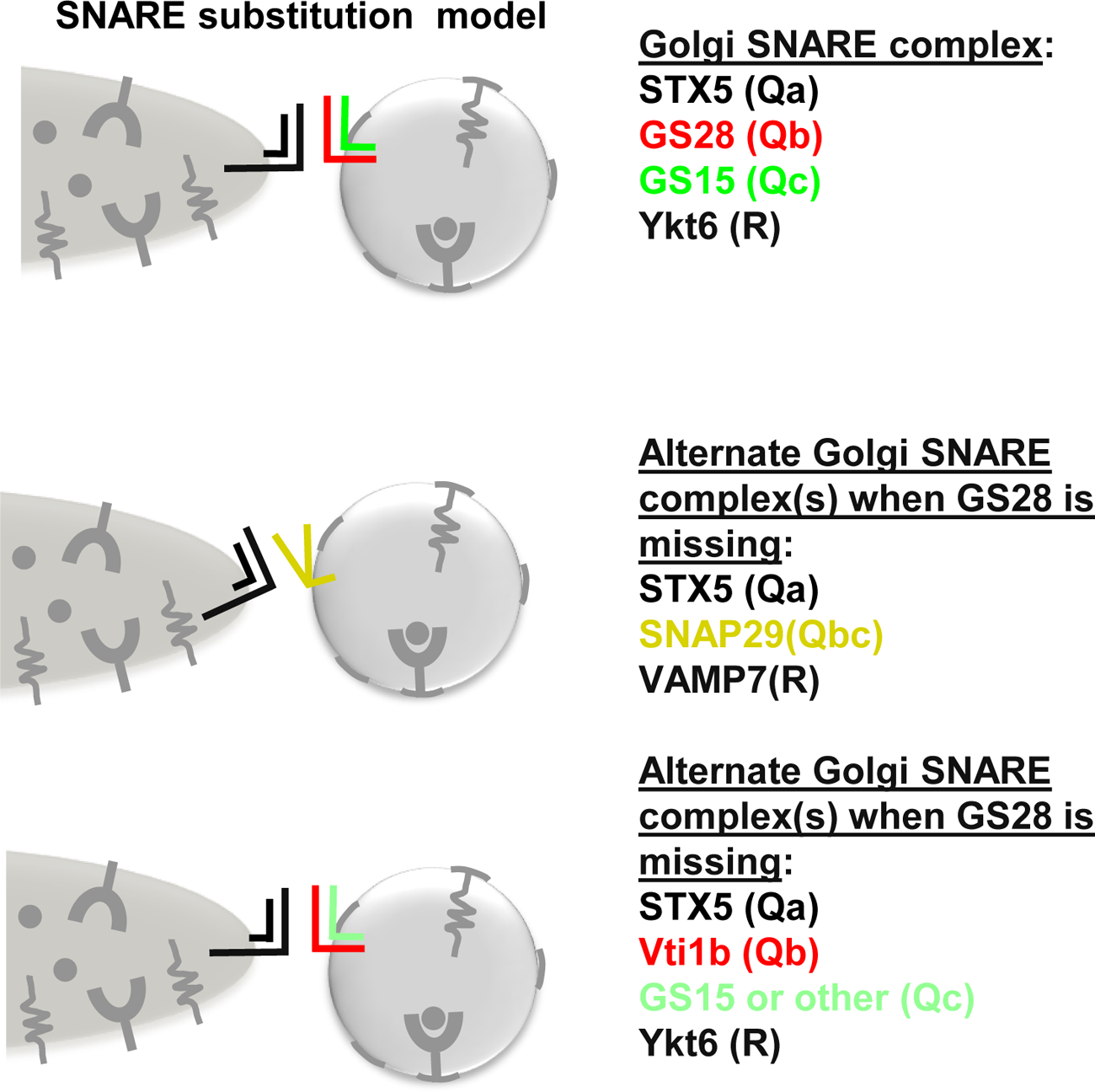
Model for novel non-canonical Golgi SNARE partnering in cells lacking GS28. SNAP29 and Vti1b are GS28 substitutes. SNAP29 operates in a complex with STX5 and VAMP7 while Vit1b could operate in a complex with Stx5, GS15 and Ykt6.

Next, we used Airyscan microscopy to analyze the localization of SNAP29 in WT and GS28 KO cells. We found a significant increase in SNAP29 colocalization with STX5 in the Golgi area in GS28 KO cells compared to WT cells, strongly supporting the model that in GS28 KO, there is an increased STX5-SNAP29 interaction at the Golgi (Figure 8). We did not detect a significant change in the colocalization between STX5 and Vti1b in GS28 deficient cells. However, Vti1b is already a perinuclear localized protein with partial colocalization with STX5 in WT cells so its increased interaction with STX5 may not lead to a visible change in Vti1b intracellular distribution (Supplemental figure 3).

**Figure 8:**
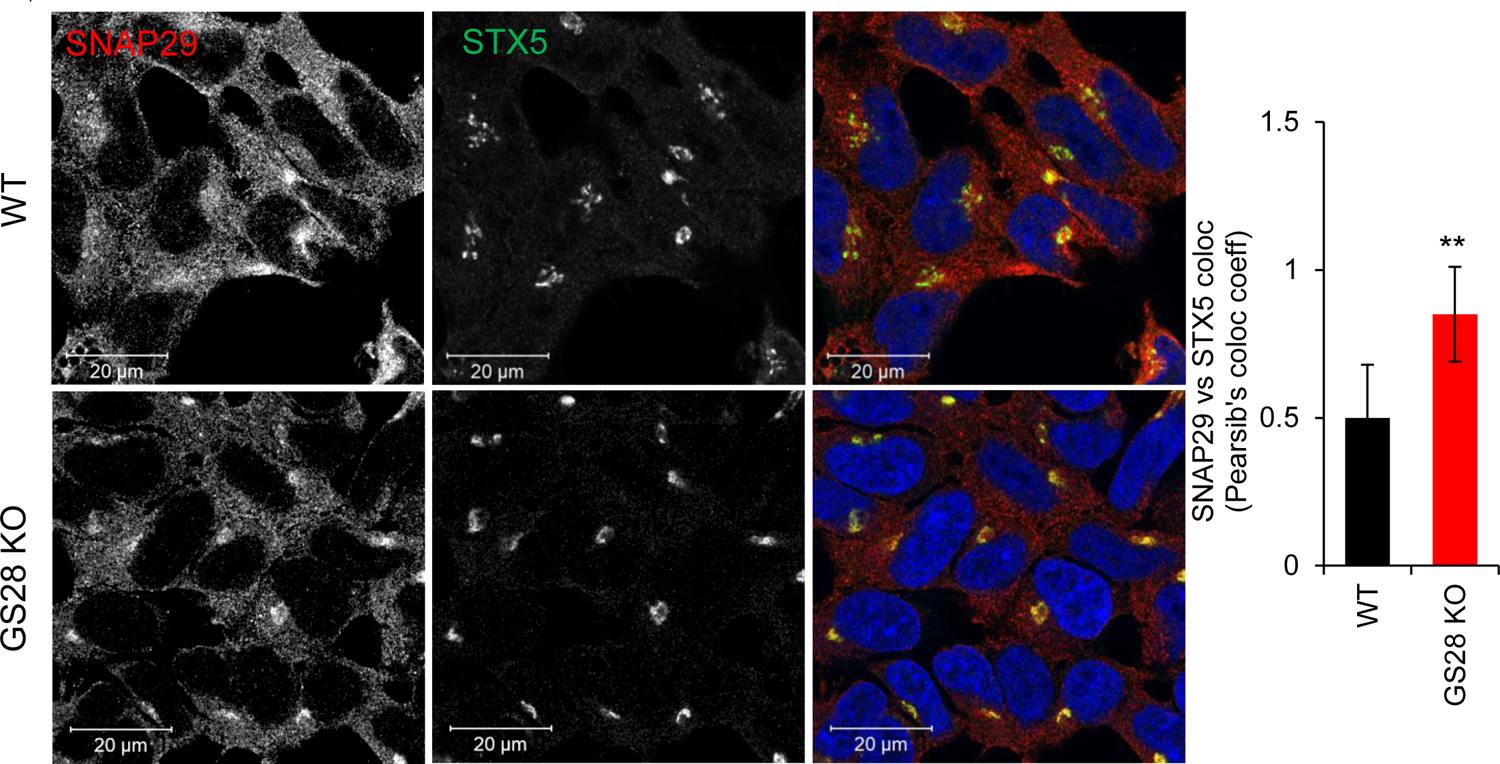
SNAP29 is significantly relocated to the Golgi upon GS28 depletion. Colocalization between STX5 and SNAP29 shows specific redistribution of SNAP29 to the Golgi region and a significant increase in SNAP29 colocalization with STX5 upon GS28 KO. Scale bars are 10µm.**p<0.001

If the SNARE substitution model is correct, we reasoned that knocking-out SNAP29 or Vti1b in GS28 KO cells will disrupt their respective non-canonical STX5-bearing SNARE complexes that in turn would further decrease intra-Golgi recycling of Golgi enzymes and glycosylation. To test this possibility GS28/Vti1b DKO and GS28/SNAP29 DKO cell lines were created. Both DKOs were viable, and they proliferated at WT rates (ZD, unpublished observation). Quantification of GNL binding to the plasma membrane of the single and double KOs by flow-cytometry, as described earlier, showed increased binding of the lectin to the surface glycoconjugates exposed by GS28-SNAP29 DKO and GS28-Vti1b DKO compared to GS28-GS15 DKO or WT (Figure 9A). This indicated that deleting either Vti1b or SNAP29 in the GS28 KO background led to an increased accumulation of under-glycosylated GNL-binding glycoproteins on the cell surface of double KO cells, supporting the involvement of both SNAP29 and Vti1b in Golgi glycosylation. Deletion of SNAP29 in GS28 KO cells had a pronounced effect on the Golgi structure as well. The transmission electron microscopy (TEM) analysis of the Golgi in mutant cells showed a greater percentage of fragmented and swollen Golgi in GS28-SNAP29 DKO compared to the WT or GS28-GS15 DKO (Figure 9B).

**Figure 9:**
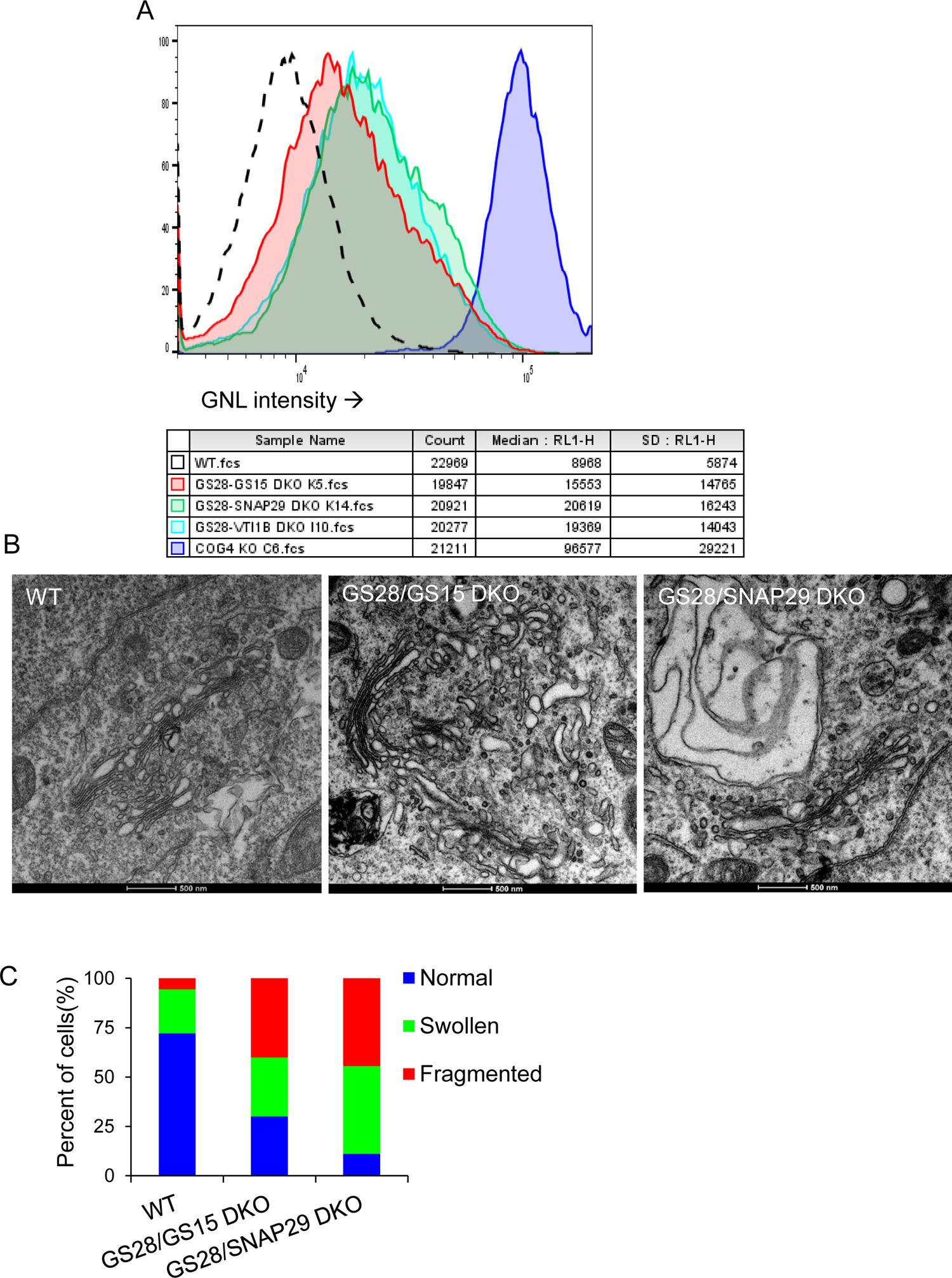
DKO of GS28/SNAP29 or GS28/Vti1b increase the severity of glycosylation defects and disrupts Golgi integrity. (A) FACS analysis revealed increase in cell-surface expression immature N-glycans is indicated by increased GNL binding to the GS29-SNAP29 or GS29-Vti1b DKOs. (B, C) EM analysis of the Golgi shows GS28-SNAP29 DKO have a higher percentage of cells with swollen and fragmented Golgi.

We have attempted to create a triple GS28/SNAP29/VTI1B KO (TKO) cell line by two complementary approaches, either by introducing VTI1B sgRNA into GS28/SNAP29 DKO cells or by introducing SNAP29 sgRNA into GS28/VTI1B DKO cells. In both cases, we were able to detect single GS28/SNAP29/VTI1B-negative cells 5-6 days after sgRNA transfection, but TKO cells did not form viable colonies, indicating that simultaneous deletion of both “canonical” and “non-canonical” STX5 partners is not tolerated in mammalian cells.

## Discussion

STX5/GS28/GS15/YKT6 is the evolutionary conserved Golgi SNARE complex that is thought to be essential for several intra-Golgi and endosome-to-Golgi vesicle trafficking steps (Banfield et al., 1995; Tai et al., 2004; Xu et al., 2002). The key Golgi vesicular tethering regulator, the COG complex, functionally and physically interacts with STX5/GS28/GS15/YKT6 assembly. COG complex defects alter Golgi SNARE complex formation and stability of both GS28 and GS15 proteins, predicting that the severe deficiency in these SNARE proteins would phenocopy defects observed in COG mutants.

To test the hypothesis, we created both single GS28 KO and GS15 KO and GS28/GS15 double KOs in HEK293T cells using the CRISPR-Cas9 strategy. Consistent with the yeast genetic data (Banfield et al., 1995; McNew et al., 1998), both GS28 and GS15 KO HEK293T cells were viable without any detectable morphological or growth defects. RPE1 cells deleted for either GS28 or GS15 were viable as well (ZD, unpublished data). GS28 deletion had a more dramatic effect on the Golgi physiology and trafficking as compared to GS15 deletion. Moreover, the deletion of GS28 caused depletion of the total levels of GS15 (Figure 2) and impaired its interaction with STX5 (Figure 8). There was also reduced Golgi localized GS15. Overall, this scenario, if not more acute, is similar to COG mutation’s/KD’s/KO’s effect on the STX5 SNARE partners (Laufman et al., 2013a; Shestakova et al., 2007; Steet and Kornfeld, 2006). Deletion of GS28 delayed retrograde trafficking of SubAB toxin to the same extent as deletion of COG subunits (Figure 3A, B) and had some impact on the stability of Golgi glycosylation machinery and Golgi glycosylation, confirming the published role of GS28 in Golgi retrograde trafficking. Surprisingly, glycosylation abnormalities in GS28 KO and GS28/GS15 DKO cells were much weaker as compared to defects observed in COG mutant cells.

There are several explanations for the limited impact of GS28/GS15 deletion on Golgi functions. One possibility is SNAREs are not essential for the maintenance and functionality of Golgi glycosylation machinery. This is unlikely because Golgi glycosyltransferases were found in v-SNARE containing Golgi-derived COPI vesicles (Adolf et al., 2019; Lin et al., 1999; Love et al., 1998) and STX5 downregulation has a dramatic effect on Golgi morphology and function (Suga et al., 2005). Another possibility is that COG regulates multiple Golgi SNARE complexes and, therefore, other Golgi-operating SNARE proteins could be more important than GS28-containing complex for the recycling and maintenance of Golgi glycosylation machinery. Indeed, the COG complex, in addition to the STX5 SNARE complex, is also shown to regulate STX16/STX6/Vti1a/VAMP4 SNARE assembly (Laufman et al., 2013a; Willett et al., 2013a), but this TGN SNARE complex has never been implicated in the recycling of Golgi enzymes, and Vti1a KO has no impact on Golgi glycosylation (ZD, unpublished data). The third possibility is that in GS28 KO cells retrograde trafficking of Golgi enzymes is rescued by cell adaptation via upregulation of an alternative non-canonical STX5-containing SNARE complex, that normally plays a minor role in Golgi biogenesis. We have tested this possibility by analyzing potential changes in STX5 partners upon deletion of GS28.

We analyzed STX5’s partner proteins by quantitative label-free MS to unbiasedly observe changes in SNARE interactions if any. Since STX5 native IPs were done under conditions that inhibited NSF activity, we reasoned that we were accumulating “fusogenic” SNARE complexes. Compared to WT, we found that in GS28 KO cells STX5’s interaction with three post-Golgi SNAREs Vti1b, VAMP7 and SNAP29 significantly increased while its interaction with ER-Golgi SNAREs GS15, BET1 and GS27 was significantly decreased in GS28 KO (Figure 6A). WB analysis of STX5 native IP was consistent with MS results (Figure 6B). STX5-Ykt6 interaction was only marginally affected by GS28 deletion and some GS15 was still Golgi-localized and bound to STX5 and Ykt6 (ZD, unpublished data), suggesting that in GS28 KO cells there might be a complex consisting of STX5/Ykt6/GS15 and Vti1b as the Qb SNARE substitute for GS28. Vti1p was first implicated in retrograde Golgi trafficking in yeasts (Lupashin et al., 1997). It was found to interact with Sed5 (Lupashin et al., 1997; von Mollard et al., 1997) and Pep12p (von Mollard et al., 1997). Dilcher et al had also found genetic interactions between Vti1 and YKT6 in yeast and proposed the existence of the Sed5p-Vt1ip-Sft1p-Ykt6p SNARE complex at the Golgi (Dilcher et al., 2001). SNARE assemblies are usually evolutionary conserved and therefore the existence of the STX5/Vti1b/GS15/Ykt6 complex is highly plausible in human cells. Unfortunately, available antibodies were not suitable for the native immunoprecipitation of Vti1b, limiting our ability to verify the exact subunit composition of STX5/Vti1b assembly. Classical in vitro studies also indicated that the positioning of Sft1 on donor liposomes was not sufficient for the fusion with Sed5-Vti1p-Ykt6 acceptor liposomes (Parlati et al., 2002). Since depletion of GS28 has not resulted in significant changes in Vti1b-STX5 colocalization, we focused our attention on STX5-SNAP29 interaction.

SNAP29 has been described as a promiscuous SNARE and its interaction with STX5, as well as many other SNAREs, has been previously reported (Hohenstein and Roche, 2001; Smeele and Vaccari, 2022; Steegmaier et al., 1998). It was also found to have various cellular distributions (Steegmaier et al., 1998) including cytosolic, perinuclear (Morelli et al., 2021; Wong et al., 1999), endolysosomal and plasma membrane possibly because of its promiscuous involvement in multiple SNARE complexes. Intracellular SNAP29 localization is flexible and could change upon changes in expression levels of its interacting partners. Schardt et al showed that overexpression of SNAP29 interactor Rab3a redistributes SNAP29 from the cytosol to the perinuclear region (Schardt et al., 2009). SNAP29, a Qbc SNARE would be the perfect substitute for deleted GS28 (Qb) and depleted GS15 (Qc). Structurally, it lacks a transmembrane domain (Steegmaier et al., 1998) allowing extra flexibility in intracellular localization. SNAP29 has been reported to form a complex with both Ykt6 and VAMP7 (Takats et al., 2018). We confirmed STX5-SNAP29 interaction at the endogenous level. Furthermore, we discovered that the endosomal R-SNARE VAMP7 but not YKT6 co-IPed with SNAP29 (Figure 6D, E Supplemental figure 3). The remarkable redistribution of SNAP29 to the Golgi bolsters our prediction that STX5/SNAP29/VAMP7 Golgi complex formation is used to compensate for the loss of STX5/GS28/GS15/YKT6 fusion machinery. To our knowledge, this is the first example of the SNARE “substitution” mechanism in mammalian cells. Previously, YKT6 was shown to substitute R-SNARE SEC22 deletion in yeast (Liu and Barlowe, 2002). Remarkably, yeast SNARE substitution also occurred for the Golgi SNARE complex containing Qa-SNARE Sed5.

Based on our experimental data, we postulated that deletion of GS28 Golgi SNARE abolished the formation of the canonical STX5 SNARE complex but did not stop retrograde trafficking in the Golgi. Vesicular trafficking in GS28 KO cells is partially compensated for by upregulation of two non-canonical SNARE complexes, STX5/SNAP29/VAMP7 and STX5/VTI1B/GS15/YKT6. To our knowledge, this is the first study to report multiple “non-canonical” SNARE complexes operating at the Golgi. We also show that distorting Golgi SNARE complexes has only marginal effects on Golgi glycosylation. The glycosylation abnormalities in COG4 KO cells are far more pronounced as compared to SNARE DKO mutants (Fig 4, 5). This surprising finding reaffirms that the COG complex is the master regulator of the Golgi.

## Materials and Methods

### List of primary antibodies and dilutions

| Antibody | Source, Catalog # | Species | WB dilution | IF dilution |
| --- | --- | --- | --- | --- |
| B4GalT1 | R&D Systems, AF-3609 | Goat | 1:500 | - |
| GALNT2 | R&D Systems, AF7507 | Sheep | 1:500 | - |
| GALNT3 | R&D Systems, AF7174 | Sheep | 1:500 | - |
| Giantin | Covance PRB- 114C | Rabbit | - | 1:1000 |
| GM130 | BD Biosciences, 610823 | Mouse | - | 1:500 |
| GS15 | Lab | Rabbit | 1:1000 | - |
| GS15 | BD Biosciences, 610961 | Mouse | - | 1:500 |
| GS28 | BD Biosciences, 611184 | Mouse | 1:500 | 1:500 |
| LAMP2 | DSHB, H4B4 | Mouse | 1:1000 | - |
| MGAT1 | Abcam, ab180578 | Rabbit | 1:500 | - |
| SDF4 | PtnTech, 10517-1-AP | Rabbit | 1:1000 | - |
| SNAP29 | Synaptic Systems, 111303 | Rabbit | 1:1000 | - |
| SNAP29 | R&D, AF7869 | Sheep | 1:1000 | 1:300 |
| SNAP29 | Abcam, ab181151 | Rabbit | 1:1000 | 1:500 |
| STX5 | Lab | Rabbit | 1:1000 | 1:1500 |
| STX5 | Santacruz, sc-365124 | Mouse | 1:500 | 1:100 |
| TGN46 | Bio-Rad, AHP500G | Sheep | 1:2000 | 1:500 |
| TMEM16 | Sigma, HPA038299 | Rabbit | 1:500 | - |
| VAMP7 | CST, 14811 | Rabbit | 1:1000 | - |
| VAMP7 | CST, 13876 | Rabbit | - | 1:300 |
| VT1IB | BD Biosciences, 611404 | Mouse | 1:250 | 1:300 |
| YKT6 | Santacruz, sc-365732 | Mouse | 1:300 | - |
| YKT6 | Bethyl, A305-479A | Rabbit | 1:500 | - |
| YKT6 | Bethyl A305-480A | Rabbit | - | 1:300 |
| β-actin | Sigma, A5441 | Mouse | 1:5000 | - |

Secondary antibodies used for WB or IF were as follows: fluorescent dye conjugated AffiniPure Donkey anti-mouse, anti-rabbit, or anti-sheep (IF 1:1000, Jackson Laboratories) and infrared dye IRDye 680RD or IRDye 800CW anti-mouse or anti-rabbit (WB 1:20,000, LI-COR).

### Cell culture

All experiments used HEK293T cells stably expressing Cas9 as described in (Khakurel et al., 2021). HEK293T-cells were cultured in DMEM /F12 (Corning) supplemented with 10% fetal bovine serum (FBS) (Gibco, Cat # 26140079, Lot #). Cells were incubated in a 37°C incubator with 5% CO2 and 90% humidity. Cells were passaged by 3min trypsinization (0.25% trypsin EDTA, Gibco) at 37°C and resuspended in media with 10% FBS. HEK293T COG4 KO cells were previously described in (Blackburn and Lupashin, 2016; D’Souza et al., 2019)

### Creation of SNARE KO stable cell lines

dual gRNAs were purchased from Transomics with the following target sequences:

GOSR1 (GS28): 1a) AAAAGAAAATATGACTTCACAGAGAGGAAT

1b) AGCGGCGGGACTCGCTCATCCTAGGGGGTG

2a) AGCCTGATCCAGAGGATCAACCTGAGGAAG

2b) AGCTTACAGGGGTAAATGATAAAATGGCAG

3a) CAGAGGATCAACCTGAGGAAGCGGCGGGAC

3b) TTTTTCTTTATTTCTAGCTTACAGGGGTAA

BET1L (GS15): 1a) AACAGAGACTCCATGGTGTTGTGCTGGACA

1b) AGACTATCATTCCGGACGTAGACGTGGCAC

2a) AGAGGATCAGAACCGGTACCTGGATGGCAT

2b) TTCTAGACCGGGAGAACAAGCGAATGGCTG

3a) CCGATTTGAGCCTGGTGACTTTGGAGGCCA

3b) CCTCTGCATCCCTATCGATGTCCAGGGCGA

Single gRNAs were purchased from Genecopia with the following target sequences SNAP29: ACAATCCGTTCGACGACGAC Vti1b: GAAGGGGTCCTCCATGGACA

Plasmids were isolated from bacteria using the QIAprep Spin Miniprep Kits (Qiagen). HEK293T-Cas9 cells were transfected using Lipofectamine 3000 (Thermo Fisher Scientific). To create GS28 KO or GS15 KO, HEK293T-Cas9 cells were transfected with a cocktail of three plasmids containing dual gRNA targets. To create GS28/GS15 DKO, GS28 KO cells were transfected with a cocktail of three plasmids containing dual gRNA targets to GS15. GS28/SNAP29 DKOs or GS28/Vti1b DKOs, GS28 KO cells were transfected with single plasmids containing one gRNA target. After 16-18h of transfection, untransfected cells were killed using 5µg/ml Puromycin for 48h. Surviving cells were then single-cell plated on 96 well plates to obtain individual colonies depleted for the target protein.

### Preparation of cell lysates and Western blotting

For the preparation of cell lysates, HEK293T WT and KO cells were grown on tissue culture dishes to a confluency of 90-100%. They were washed thrice with PBS and lysed in 2% hot SDS, heated for 10 min at 70°C. The total protein concentration was measured using the BCA protein assay (Pierce). Samples were prepared so that they were at a concentration of 20-30µg and 6X Laemmli sample buffer containing 5% β-mercaptoethanol (Sigma) was added. Samples were further heated at 70°C for 10 min. Bio-Rad (4–15%) gradient gels were used for gel electrophoresis Proteins were blotted onto 0.2 µm nitrocellulose membranes (Amersham Protran, GE Healthcare) using the Thermo Scientific Pierce G2 Fast Blotter. Membranes were rinsed in PBS, blocked in EveryBolt blocking buffer (BioRAD, Cat # 12010020) for 30 min, and incubated with primary antibodies overnight at 4°C. Membranes were washed with PBS and incubated with secondary fluorescently tagged antibodies diluted in BioRAD blocking buffer for 60 min. All the primary and secondary antibodies are listed above. Blots were then washed and imaged using the Odyssey Imaging System. Images were processed using the LI-COR Image Studio software.

### Lectin staining

Gel electrophoresis and blotting were performed as described above. After transfer, the membrane was blocked with 3% bovine serum albumin (BSA) for 30 min. The lectins HPA (Invitrogen, Cat # L32454) or GNL (Khakurel et al., 2021) conjugated to Alexa 647 fluorophore were diluted 1:1000 in 3% BSA from their stock concentration of 1 and 5 µg/µl, respectively. Blots were incubated with lectin solutions for 30 min and then washed in PBS four times for 4 min each and imaged using the Odyssey Imaging System.

### Flow Cytometry

Cells were grown to 80–90% confluency on a 12-well plate. On the day of the experiment, cells were detached by incubating with 10 mM EDTA in 1X PBS at 37°C. After three PBS washes, to get rid of the EDTA, cells were resuspended in ice-cold 0.1% BSA. The lectins used were GNL (Khakurel et al., 2021), HPA (Invitrogen, Cat #L32454), CTX (Invitrogen, Cat # C34776). They were diluted at 1:500 in ice-cold 0.1% BSA and incubated with the cells for 30 min on ice. DAPI was added to the samples just prior to the analysis and samples were run on the Attune NxT flow cytometer (Thermo Fisher Scientific). Dead cells were excluded by gating on live (DAPI^-^) cells. Singlets were gated forward scatter height (FSC) vs FSC area density plots. Final gating was done in a side scatter area vs FCS density plot. The median fluorescence (of the lectin’s fluorophore) intensity for this population was obtained from a histogram plot.

### Immunoprecipitation assay

WT and KO cells were plated on 15 cm dishes. Cells were grown to 100% confluency. Prior to lysis, cells were treated with 1mM NEM for 15min at 37°C in order to inhibit NSF activity and thereby preserve SNARE complexes (Laufman et al., 2013a; Mallard et al., 2002; Perez-Victoria and Bonifacino, 2009). Thereafter, cells were washed thrice with DPBS and lysed for 1h on ice in 1% Triton lysis buffer made in 50 mM Tris pH 7.5, 150 mM NaCl, 0.1 mM PMSF and 1X Protease inhibitor cocktail. Lysates were centrifuged at 20,000 r.c.f for 10 min at 4°C. The supernatant was collected. A small volume of the supernatant was set aside as the “input”. For STX5 IP, STX5 (Lab) antibody was added to the remaining supernatant at a 1:100 dilution. For SNAP29 IP, (Abcam) was added at a 1:100 dilution. Lysates were incubated with the antibody at 4°C, for 16h on a rotor. Antigen-antibodies complexes were pulled down using Protein G-Agarose (Roche, Lot #70470320). First, the beads were washed thrice in the wash buffer composed of 0.05% Triton in the Tris-NaCl buffer described above and then introduced into the lysates. The beads were incubated with the lysates on a rotor for 90 min at RT. The beads were collected by centrifugation and washed trice in the 0.05% Triton wash buffer. Bound protein was eluted by heating the beads in 2X Laemmli sample buffer containing 10% β-mercaptoethanol at 95°C for 10 min. Western blot analysis for the IP samples was performed as described above.

### Mass-spectrometry of STX5 IP and data analysis

The STX5 IP was performed as described above. For MS, before eluting, the Protein-G Agarose beads were washed three more times in 1X PBS to get rid of the excess detergent. Elution was performed as described above. Sample preparation for MS was described in (Wisniewski et al., 2009). Briefly, purified proteins were reduced, alkylated, and digested using filter-aided sample preparation. Tryptic peptides were then separated by reverse-phase XSelect CSH C18 2.5 um resin (Waters) on an in-line 150 x 0.075 mm column using an UltiMate 3000 RSLCnano system (Thermo). Peptides were eluted using a 60 min gradient from 98:2 to 65:35 buffer A:B ratio. Eluted peptides were ionized by electrospray (2.2 kV) followed by mass spectrometric analysis on an Orbitrap Exploris 480 mass spectrometer (Thermo). To assemble a chromatogram library, six gas-phase fractions were acquired on the Orbitrap Exploris with 4 m/z DIA spectra (4 m/z precursor isolation windows at 30,000 resolution, normalized AGC target 100%, maximum inject time 66 ms) using a staggered window pattern from narrow mass ranges using optimized window placements. Precursor spectra were acquired after each DIA duty cycle, spanning the m/z range of the gas-phase fraction (i.e. 496-602 m/z, 60,000 resolution, normalized AGC target 100%, maximum injection time 50 ms). For wide-window acquisitions, the Orbitrap Exploris was configured to acquire a precursor scan (385-1015 m/z, 60,000 resolution, normalized AGC target 100%, maximum injection time 50 ms) followed by 50x 12 m/z DIA spectra (12 m/z precursor isolation windows at 15,000 resolution, normalized AGC target 100%, maximum injection time 33 ms) using a staggered window pattern with optimized window placements. Precursor spectra were acquired after each DIA duty cycle.

Following data acquisition, spectra were searched using an empirically corrected library and a quantitative analysis was performed to obtain a comprehensive proteomic profile. Proteins will be identified and quantified using EncyclopeDIA (Searle et al., 2018). Scaffold DIA was used for visualization. False discovery rate (FDR) thresholds, at both the protein and peptide levels, were set at 1%. Protein exclusive intensity values were assessed for quality using ProteiNorm (Graw et al., 2020). Popular normalization methods were evaluated including log2 normalization (Log2), median normalization (Median), mean normalization (Mean), variance stabilizing normalization (VSN) (Huber et al., 2002), quantile normalization (Quantile) (Bolstad, 2022) cyclic loess normalization (Cyclic Loess) (Ritchie et al., 2015), global robust linear regression normalization (RLR) (Chawade et al., 2014), and global intensity normalization (Global Intensity) (Chawade et al., 2014). The individual performance of each method was evaluated by comparing the following metrices: total intensity, Pooled intragroup Coefficient of Variation (PCV), Pooled intragroup Median Absolute Deviation (PMAD), Pooled intragroup estimate of variance (PEV), intragroup correlation, sample correlation heatmap (Pearson), and log2-ratio distributions. The VSN normalized data was used to perform statistical analysis using Linear Models for Microarray Data (limma) with empirical Bayes (eBayes) smoothing to the standard errors (Ritchie et al., 2015). Proteins with an FDR adjusted p-value ≤0.05 and a fold change ≥2 were considered significant.

### Immunofluorescence (IF) microscopy

12 mm glass coverslips (#1, 0.17 mm thickness) were collagen-coated. WT and KO cells were plated to be 60–70% confluent at the time of processing. Cells were washed with Dulbecco’s phosphate-buffered saline (DPBS) and stained according to the protocol described previously (Zolov and Lupashin, 2005). Briefly, freshly prepared 4% paraformaldehyde (PFA) (16% stock solution diluted in DPBS; Electron Microscopy Sciences) was used as a fixative. After fixation, cells were permeabilized with 0.1% Triton X-100 for 1 min followed by treatment with 50 mM ammonium chloride for 5 min to quench free aldehydes. Cells were then washed three times with DPBS and blocked twice for 10 min each with 1% BSA, and 0.1% saponin in DPBS. Antibodies were diluted in DPBS with 1% cold fish gelatin and 0.1% saponin. Cells were incubated with the primary antibody for 1 h at room temperature. Cells were washed four times with DPBS and incubated for 30 min with fluorescently tagged secondary antibodies diluted in antibody buffer. Cells were washed four times with DPBS. Hoechst diluted 1:10000 in DPBS was used to stain the nucleus. Coverslips were then washed four times with DPBS, rinsed with ddH2O, and mounted on glass microscope slides using Prolong Gold anti-fade reagent (Life Technologies).

In the case of staining the plasma membrane with GNL lectin, a step fixation method was followed. In the first set, cells on coverslips were fixed with 1% PFA for 15min, followed by DPBS washes. 1% BSA in DPBS was used for blocking. After 30min blocking cells were incubated with the lectin GNL, diluted to 1:500 in 1% BSA blocking buffer. Coverslips were washed thrice with PBS and the above described IF protocol was continued.

Cells were imaged with the 63X oil 1.4 numerical aperture (NA) objective of the LSM880 using the Airyscan module. Pearson’s colocalization analysis was performed in ZenBlue version 2.6. Images were converted to TIF format and assembled using Adobe Photoshop CS6.

### Transmission Electron Microscopy (TEM)

Samples were processed for TEM according to Valdivia’s lab protocol (Cocchiaro and Valdivia, 2009) with some modifications (Pokrovskaya et al., 2012). Briefly, excess growth media was removed from the dish with cells and an equal volume of 1X fixative was added for 5 min at RT effectively bringing the fixative to 0.5X concentration. 0.5X fixative was then replaced with 1X fixative for 10 min at RT. 1X fixative was composed of 4% PFA (EMS) and 1% GA (EMS) in 0.1M Phosphate buffer, pH 7.4. Finally, cells were fixed for 20 min on ice with 2.5% GA and 0.05% malachite green (EMS) in 0.1 M sodium cacodylate buffer, pH 6.8. Cells were washed 4 x 5min with 0.1M sodium cacodylate buffer and post-fixed for 30 min at RT with 0.5% osmium tetroxide and 0.8% potassium ferricyanide in 0.1 M sodium cacodylate buffer, washed again and incubated for 20 min on ice in 1% tannic acid. After washing in buffer and H2O samples were incubated for 1 h in 1% uranyl acetate at RT. Specimens were gradually dehydrated in a graded series of increasing ethanol concentrations, washed with Propylene Oxide (EMS) and incubated in 50% PO/resin mixture before embedding in Araldite 502/Embed 812 resins (EMS). Ultrathin sections were imaged at 80 kV on the FEI Technai G2 TF20 transmission electron microscope. Digital images were acquired with FEI Eagle 4kX USB Digital Camera.

## Statistical analysis

All results are representatives of at least three biological replicates. WBs were quantified by densitometry using the LI-COR Image Studio software. For statistical analysis, a student’s t-test was performed in Microsoft Excel 2010. All graphs were plotted in Microsoft Excel 2010. Error bars represent standard deviation.

## Supporting information

STX5 IP mass-spectrometry data

## Author Contributions

Z.D. wrote the article and made substantial contributions to conception and design, acquisition of data, analysis, and interpretation of data. I.D.P. performed EM experiments and interpreted the data. V.L. edited the article and made substantial contributions to conception and design.

## Acknowledgements

We are thankful to all our colleagues who provided reagents. We are thankful to Tetyana Kudlyk for excellent technical support as well as Farhana Taher Sumya and Amrita Khakurel for critical discussion. We would also like to thank the UAMS Digital Microscopy and Proteomics cores for the use of their facilities and expertise.

---

**Supplemental figure 1: GS15 immunoprecipitation in WT and GS28 KO cells shows impaired SNARE interactions in GS28 KO**. CoIP of GS15 partners using affinity purified polyclonal antibodies (Lab made) to GS15 shows a marked reduction in the level of SNARE partners, STX5 and Ykt6 coIPed in GS28 KO compared to WT.

**Supplemental figure 2: Ykt6 is not the R-SNARE partner of SNAP29.** CoIP of SNAP29 partners using affinity purified antibodies (Abcam) to SNAP29 shows that this SNARE does not interact with COG3 or Ykt6.

**Supplemental figure 3: Co-staining WT and GS28 KO cells with STX5 and Vti1b or shows Vti1b has a peri-nuclear distribution and colocalizes with STX5.** WT and GS28 KO cells were treated with 1mM NEM for 15min at 37°C prior to staining. Vti1b is a perinuclear SNARE and colocalizes with STX5. There was no significant change in colocalization between STX5 and Vti1b in WT and GS28 KO. n≥30

**Supplemental table 1: Normalized DIA MS data of STX5 IP in WT vs GS28 KO.** Fold change (log_2_ FC) and significance (p value) of all STX5 interactors in WT vs GS28 KO.

