## Supplemental Figure 1 for "STX5’s flexibility in SNARE pairing supports Golgi functions"

**S1**

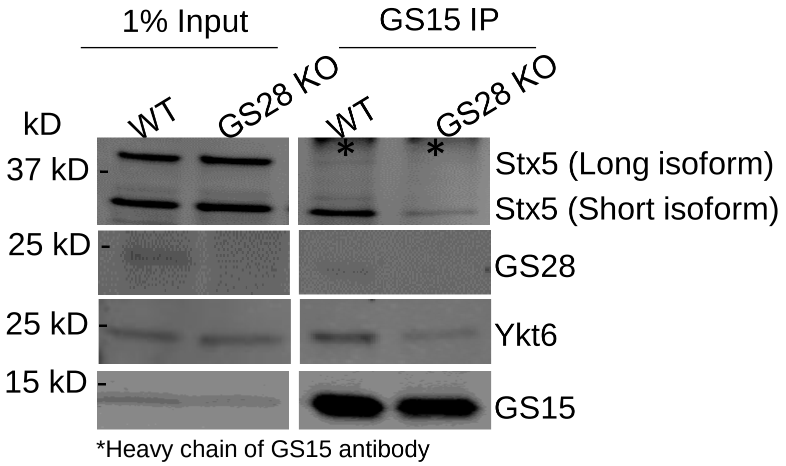


**Supplemental figure 1: GS15 immunoprecipitation in WT and GS28 KO cells shows impaired SNARE interactions in GS28 KO.** CoIP of GS15 partners using affinity purified polyclonal antibodies (Lab made) to GS15 shows a marked reduction in the level of SNARE partners, STX5 and Ykt6 coIPed in GS28 KO compared to WT.
