## Supplemental Figure 2 for "STX5’s flexibility in SNARE pairing supports Golgi functions"

**S2**

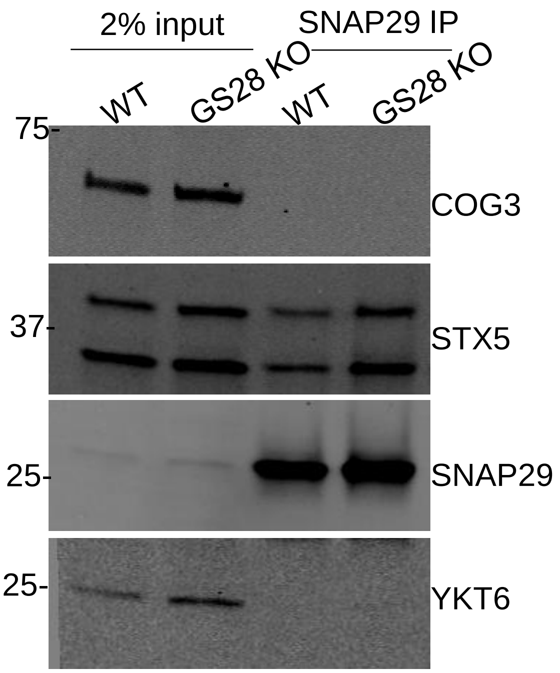


**Supplemental figure 2: Ykt6 is not the R-SNARE partner of SNAP29.** CoIP of SNAP29 partners using affinity purified antibodies (Abcam) to SNAP29 shows that this SNARE does not interact with COG3 or Ykt6.
