## Supplemental Figure 3 for "STX5’s flexibility in SNARE pairing supports Golgi functions"

**S3**

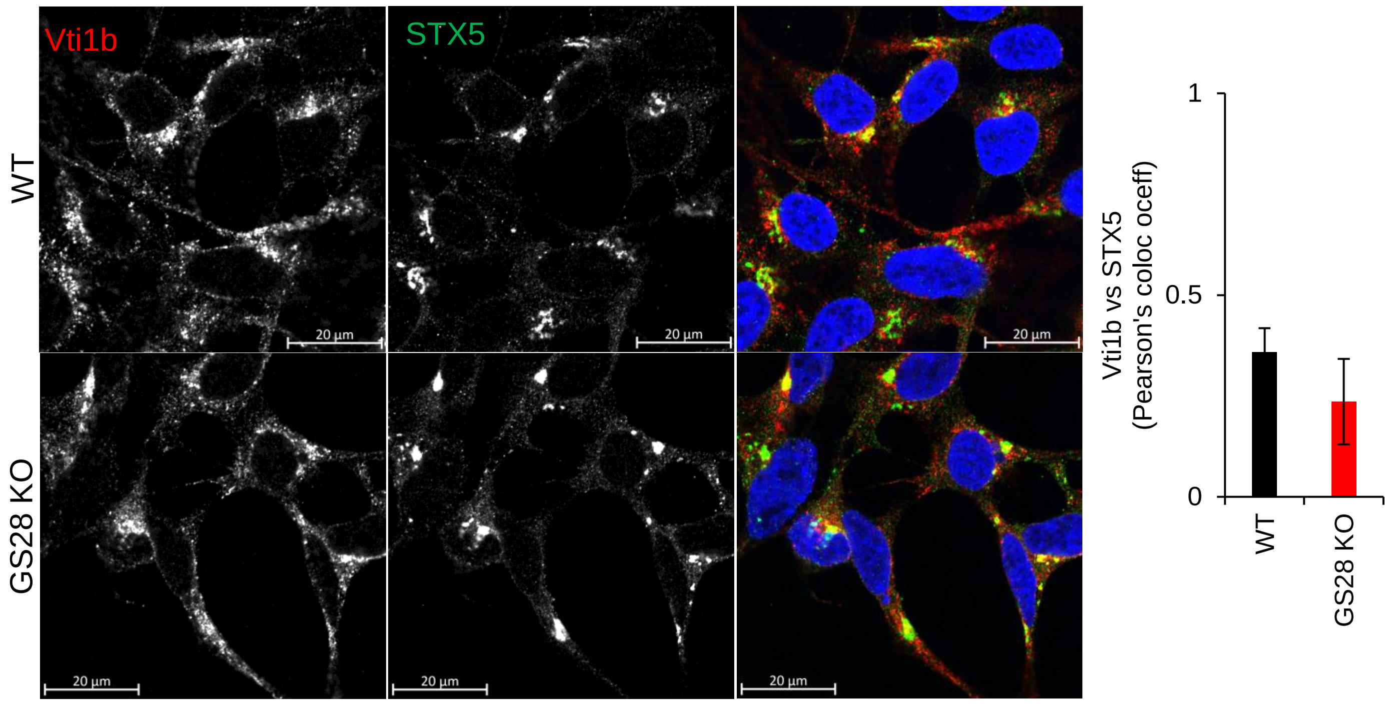


**Supplemental figure 3: Co-staining WT and GS28 KO cells with STX5 and Vti1b or shows these Vti1b has a peri-nuclear distribution and colocalizes with STX5.** WT and GS28 KO cells were treated with 1mM NEM for 15min at 37°C prior to staining. Vti1b is a perinuclear SNARE and colocalizes with STX5. There was no significant change in colocalization between STX5 and Vti1b in WT and GS28 KO. n≥30
